# Threat-induced hippocampal connectivity fingerprints do not generalize to psychosocial stress

**DOI:** 10.1101/2020.11.16.380220

**Authors:** Anne Kühnel, Michael Czisch, Philipp G. Sämann, BeCOME study team, Elisabeth B. Binder, Nils B. Kroemer

## Abstract

Stress is an everyday experience and maladaptive responses play a crucial role in the etiology of affective disorders. Despite its ubiquity, the neural underpinnings of subjective stress experiences have not yet been elucidated, particularly at an individual level. In an important advance, Goldfarb et al.^1^ showed recently that subjective stress and arousal levels in response to threatening stimuli were successfully predicted based on changes in hippocampal connectivity during the task using a machine learning approach. Crucially, stress responses were predicted by interpretable hippocampal connectivity networks, shedding new light on the role of the hippocampus in regulating stress reactivity^2^. However, the authors induced stress by displaying aversive pictures, while stress research often relies on the extensively validated Trier social stress task (TSST)^3^. The TSST incorporates crucial factors such as unpredictability of success and the social-evaluative threat of the stressor thereby eliciting cortisol responses more robustly compared to threatening images^4^. Towards generalization, cross validation within a sample as conducted by Goldfarb et al.^1^ or independent replications are important steps, but the generalizability to different stressors allows to draw broader conclusions about the potential use of hippocampal connectivity to predict subjective stress^5^. Arguably, translating these findings to clinical applications would require a broad generalization of the results or the prediction algorithm to psychosocial stress. Here, we assessed the predictive performance of Goldfarb et al’s^1^ algorithm for subjective stress in an independent sample using an MR adaption of the TSST^6,7^. In line with Goldfarb et al.^1^, we observed robust stress-induced changes in hippocampal connectivity. However, the spatial correlation of the changes in connectivity was low indicating little convergence across alleged stress paradigms. Critically, stress-induced changes of hippocampal connectivity were not robustly predictive of subjective stress across a multiverse of analyses based on connectivity changes. Collectively, this indicates that the generalizability of the reported stress connectivity fingerprint to other stressors is limited at best, suggesting that specific tasks might require tailored algorithms to robustly predict stress above chance levels.

## Methods

To evaluate generalization, we derived complementary operationalizations of stress-induced changes in hippocampal-seed connectivity using previously reported data from an adaptation of the TSST for neuroimaging^8^ (N=67)^7^. First, we applied the emulated the method to estimate background functional connectivity as closely as possible in psychosocial stress and calculated individual changes in hippocampal connectivity during Stress compared to PreStress (Figure 1a). Second, as predictive performance was limited, we used generalized psychophysiological interactions (gPPI^9^). gPPI is commonly applied to study task-related functional connectivity and estimates task-induced activity and connectivity changes for multiple conditions and phases simultaneously, instead of calculating functional connectivity for each phase separately. Third, we applied linear mixed-effects models predicting extracted time series of the complete task while including a preceding resting-state measurement as a stable baseline. To predict previously reported^7^ changes in positive or negative affect immediately and 30 minutes after the task, we applied the same machine learning algorithm as reported^1,10^. To assess within-sample generalization, we applied leave-one-out (LOOCV) and 10fold cross validation (10foldCV). In line with Goldfarb et al.^1^, we performed additional permutation tests for the 10foldCV models. However, instead of comparing the null distribution with a parameter distribution across 1,000 cross-validation sets using a Kolmogorov-Smirnow test, we calculated permutation p-values by comparing the observed correlation with the null-distribution leading to more conservative significance estimates.

**Figure 1:**
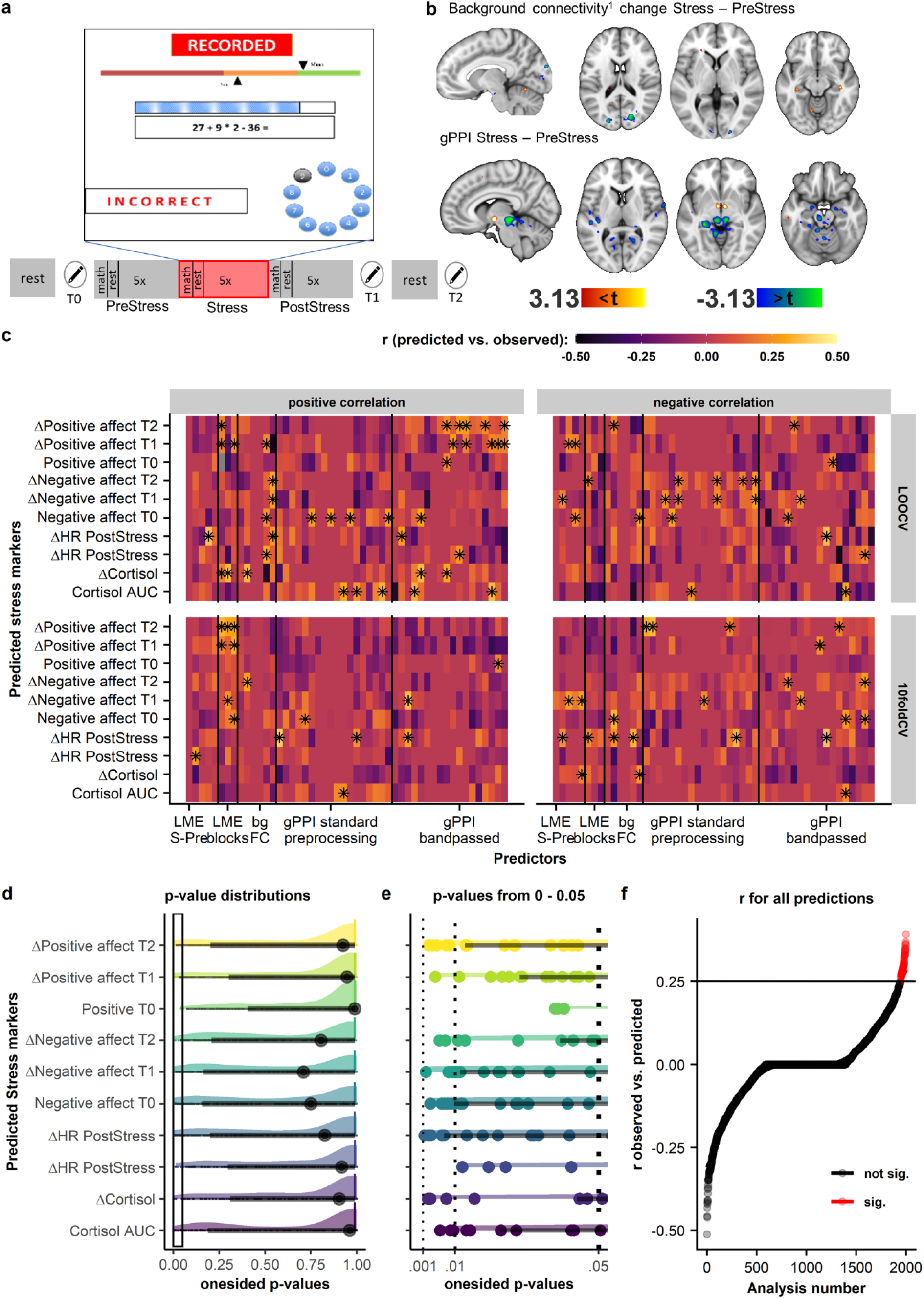
Changes in hippocampal connectivity do not robustly predict subjective or physiological stress markers. a) Task design of the psychosocial stress task including an example screen shown during the stress phase. Each phase consists of 5, 50s blocks of math problems interleaved with 40s rest/baseline blocks. b) Whole-brain changes of bilateral hippocampal connectivity during Stress compared to a neutral PreStress condition (same math problems, no aversive elements). The upper panel shows changes in background connectivity estimated as in Goldfarb et al.^1^. The lower panel shows the group-level effect of the gPPI contrast Stress-PreStress showing increasing connectivity to the hypothalamus and decreasing connectivity to the posterior midbrain. Voxel-threshold p < .001, uncorrected. c) Correlations between observed vs. predicted stress markers for all (500×2 (positive vs. negative) × 2 (LOOCV vs. 10foldCV)) models. Significant (one-sided p < .05, not corrected for multiple testing) predictions are marked with asterisks. Predictions were not robust across leave-one-out cross validation (LOOCV) and 10fold cross-validation (10fold) or different operationalizations of stress effects. The first models include time series based linear mixed-effects (LME) estimating either Stress vs. PreStress (LME S-Pre) or task-induced changes vs. baseline/rest (LME blocks) and include connectivities within a pre-defined network or between the hippocampus and 45 ROIs defined based on peak coordinates from Goldfarb et al.^1^. In the second model (bg FC) stress-induced changes in background hippocampal functional connectivity were estimated emulating the method in Goldfarb. The third set of models was based on whole-brain generalized psychophysiological interaction (gPPI) analyses with a hippocampus seed, either using standard preprocessing or including additional bandpass filtering (gPPI bandpassed). For all gPPIs, predictors were individual level estimates from the contrast Stress – PreStress or PostStress – PreStress extracted from a functional atlas, the main effect clusters, or the main effect clusters reported in Goldfarb et al.^1^ All hippocampus-specific analyses were also repeated for the anterior and posterior hippocampus^11^ separately. d) The distribution of one-sided p-values across all models and predicted indices has its peak close to p = 1 and no prominent peaks at p < .05. e) The same p-value distributions zoomed in in the interval between 0 and 0.05 show only a small number of significant (p < .05) models and none with a p < .001. f) Correlation coefficients (r) of the correlation predicted vs. observed from all 100 × 2 × 10 models showing only 5.05% of significant (marked in red) models (one-sided p < .05, N = 101).

## Results

To assess whole-brain changes in hippocampus-seed connectivity in the psychosocial stress task, we used the same analysis pipeline as Goldfarb et al.^1^ We preprocessed our data as previously reported and regressed out movement, global signal and task effects^7^. We calculated changes in z-transformed hippocampal connectivity for each task block compared to its preceding baseline, and averaged changes across blocks of the same condition for prediction. During stress, hippocampal connectivity to the right cuneus and left middle occipital gyrus decreased compared to the neutral PreStress condition. However, changes in connectivity showed very limited spatial correlation with the reported effects in Goldfarb et al.^1^ (voxel-wise whole-brain r= .005, atlas ROIs^11^ whole-brain r= .16, voxel-wise within different networks^12^ rs: -.07 - .03) on the whole-brain level. We extracted individual estimates for prediction using three sets of predictors: first from a connectivity-based, whole-brain atlas (268 ROIs^11^), second from significant group-level clusters (p_voxel_ < .001, p_cluster.uncorrected_ < .05), and third from the main effect ROIs (entire hippocampus) reported in Goldfarb et al.^1^. Nonetheless, none of the subjective stress markers could be robustly (i.e. using LOOCV and 10foldCV) predicted (best: ΔT2 Negative LOOCV: r= .31, p= .005 corresponding 10fold: r= .01, p= .45). Likewise, a more direct replication using the clusters identified by Goldfarb et al.^1^ did not robustly predict subjective stress in our psychosocial stress paradigm (Table 1).

**Table 1:** Predictive performance of changes in hippocampal connectivity during stress (background connectivity^1^) using the same clusters as Goldfarb et al.^1^

|  | LOOCV |  | 10foldCV |  |  |  |
| --- | --- | --- | --- | --- | --- | --- |
|  | r | p | r | p | p<sub>perm</sub> | p<sub>perm</sub> KS |
| ΔPositive T1 (+) | .00 | .99 | .00 | .99 | 1 | < .001 |
| ΔNegative T1 (+) | .11 | .17 | .08 | .25 | .23 | < .001 |
| ΔPositive T1 (-) | .12 | .15 | -.30 | .99 | .99 | < .001 |
| ΔNegative T1 (-) | -.20 | .95 | .13 | .14 | .13 | < .001 |

*Note:* r refers to Pearson correlation coefficients, in case of the 10fold cross validation the median r across 1,000 iterations with different splits is reported. P values are one-sided (positive correlation), p<sub>perm</sub> is the p value corresponding to a permutation test. Here, the null distribution is estimated by 1,000 repetitions of the prediction algorithm after random resamples of the target variable. The p-value reflects the rank of the observed r compared to the null distribution. P<sub>perm</sub> KS refers to the p-value derived by comparing the null distribution to a distribution of the observed value across 1,000 different cross-validation splits (as described in Goldfarb et al.<sup>1</sup>) using a Kolmogorov-Smirnov test.

Since this method only showed significant changes in hippocampal connectivity during stress to two occipital clusters, we tested if using gPPI (i.e., the common method to estimate task-induced changes in seed-based functional connectivity), either with or without bandpass filtering (Figure 1b) and global signal regression would improve predictions. Here, first-level models^7^ additionally included the bilateral hippocampal time course and interactions for each phase (Stress, PostStress; PreStress as reference). Again, none of the stress markers could be robustly predicted using both LOOCV and 10foldCV based on comparisons of Stress and PreStress (best: ΔT1 Negative LOOCV: r= .29, p= .009 corresponding 10fold: r= -.11, p= .82). Intriguingly, predictors derived from changes in hippocampal connectivity during PostStress predicted changes in negative affect (LOOCV: r= .25, p= .02, corresponding 10fold: r= .28, p= .01), but this phase was not available in the study by Goldfarb et al.^1^.

Subsequently, we explored if incorporating a preceding resting-state baseline and estimating stress-induced connectivity changes using linear mixed-effects models based on detrended, denoised, concatenated data would lead to an improved prediction. We estimated connectivity changes using pairwise connectivities between hippocampus seeds and atlas ROIs^11^ corresponding to the peak coordinates of the reported main effects^1^. Comparable to gPPI, the models (one for each pairwise connectivity) included regressors for each task block, the seed time series, and their interactions, all as random effects. In line with previous analyses, hippocampal connectivity changes did not robustly predict stress-induced effects with LOOCV and 10fold CV (best: ΔT1 Positive LOOCV r= .23, p= .028, corresponding 10fold: r= -.05, p= .66). Instead, hippocampal connectivity changes predicted changes in positive affect (LOOCV r= .22 p= .035, 10fold r= .22, p= .04), but this was driven by changes between and rest and the PostStress phase of the task.

To assess the prediction of the proposed algorithm for physiological stress responses, we also predicted the previously reported^7^ biomarkers of stress, increase in heart rate and cortisol response, using the same procedures. While we observed few (3) successful predictions across LOOCV and 10fold CV (Figure 1c), they were not robust across different operationalizations such as slight changes in preprocessing or varying definitions of ROIs indicating limited generalizability. Nonetheless, other, more fine-grained operationalizations of the physiological response may be more robustly predicted by task-induced changes in neural connectivity.

## Discussion

To summarize, in contrast to a task showing aversive visual stimuli, stress-induced connectivity changes in the hippocampus did not robustly predict subjective or physiological markers of stress in an established psychosocial stress task. While some models predicted stress responses above chance level (6%) using LOOCV, only 7 out of a multiverse of 1,000 models were also predictive after 10foldCV, which has been recommended to guard against overfitting^13^. A lack of replicability and convergence of reported neural underpinnings of stress and the associated subjective or physiological responses has been an ongoing matter of debate^14^. These differences may be due to common issues in neuroimaging^15^ such as small sample sizes, flexible analyses methods^16^, or publication bias. However, as detailed by Yarkoni^5^, those explanations neglect the issue of generalizability beyond the strict specifications of a single experiment to a broader understanding of an underlying mechanism. This lack of generalizability could explain why no converging stress fingerprints or even correlates have been identified yet. Stress paradigms and experimental procedures vary widely^17^ and may not even elicit comparable changes in functional connectivity. This is not only an issue in neuroimaging research, but also applies more generally to investigations of endocrine or other physiological stress effects. It has already been shown that even slight modifications of procedures surrounding a stress task can affect individual stress reactions^18^, further highlighting the difficulties of generalizing effects across different settings allegedly testing the same concept. Discovering connectivity profiles related to stress responses remains critical to understand not only adaptive stress reactivity, but maladaptive stress responses in mental disorders. However, our results highlight that caution is warranted in drawing broad conceptual conclusions before evaluating the predictive performance across different operationalization of the same construct. In light of our findings, it is plausible that viewing aversive pictures versus failing arithmetic problems in front of an audience involve different kinds of stress responses that may be regulated by different mechanisms.

## Acknowledgement

NBK received salary support from the University of Tübingen, fortune grant #2453-0-0.

## Author contributions

EBB, PGS, MC and NBK were responsible for the study concept and design. AK & NBK conceived the method. AK performed the data analysis and NBK contributed to analyses. AK & NBK wrote the manuscript. All authors contributed to the interpretation of findings, provided critical revision of the manuscript for important intellectual content and approved the final version for publication.

